# Sir2 mitigates an intrinsic imbalance in origin licensing efficiency between early and late replicating euchromatin

**DOI:** 10.1101/2020.01.24.918862

**Authors:** Timothy Hoggard, Carolin A. Müller, Conrad A. Nieduszynski, Michael Weinreich, Catherine A. Fox

## Abstract

A eukaryotic chromosome relies on the function of multiple spatially distributed DNA replication origins for its stable inheritance. The location of an origin is determined by the chromosomal position of an MCM complex, the inactive form of the DNA replicative helicase that is assembled on chromosomal DNA in G1-phase (a.k.a. origin licensing). While the biochemistry of origin licensing is understood, the mechanisms that promote an adequate spatial distribution of MCM complexes across chromosomes are not. We have elucidated a role for the Sir2 histone deacetylase in establishing the normal distribution of MCM complexes across *Saccharomyces cerevisiae* chromosomes. In the absence of Sir2, MCM complexes accumulated within both early-replicating euchromatin and telomeric heterochromatin, and replication activity within these regions was enhanced. Concomitantly, the duplication of several regions of late-replicating euchromatin were delayed. Thus, Sir2-mediated attenuation of origin licensing established the normal spatial distribution of origins across yeast chromosomes required for normal genome duplication.

**Significance statement:** In eukaryotes, multiple DNA replication origins, the sites where new DNA synthesis begins during the process of cell division, must be adequately distributed across chromosomes to maintain normal cell proliferation and genome stability. This study describes a repressive chromatin-mediated mechanism that acts at the level of individual origins to attenuate the efficiency of origin formation. This attenuation is essential for achieving the normal spatial distribution of origins across the chromosomes of the eukaryotic microbe *Saccharomyces cerevisiae*. While the importance of chromosomal origin distribution to cellular fitness is now widely acknowledged, this study is the first to define a specific chromatin modification that establishes the normal spatial distribution of origins across a eukaryotic genome.

## Introduction

An adequate distribution of DNA replication origins across eukaryotic chromosomes is important for maintaining cell proliferation and genome stability over the course of multiple rounds of cell division. Chromosomal regions that contain a paucity of origins are linked to chromosome fragility and cancer-associated deletions (1; 2; 3; 4). Origin function relies on two distinct cell-cycle restricted reactions (5). In G1-phase, the ORC (**o**rigin **r**ecognition **c**omplex) and Cdc6 protein bind DNA and direct the assembly of a stable catalytically inactive MCM (**m**ini**c**hromosome **m**aintenance) complex, a.k.a. origin licensing. In S-phase, kinases and accessory proteins act on the MCM complex to convert it into two bidirectionally oriented replicative helicases that unwind the DNA to allow for new DNA synthesis (a.k.a. origin activation). Thus, the effective chromosomal distribution of two distinct reactions, MCM complex loading and MCM complex activation, establishes the normal spatiotemporal distribution of origins. While recent progress in *Sacccharomyces cerevisiae* has provided insights into mechanisms that regulate the distribution of the S-phase MCM complex activation reaction, little is known about how the chromosomal distribution of MCM complexes is achieved (6; 7; 8; 9; 10; 11).

Several attributes of *S. cerevisiae* are helpful for addressing mechanisms relevant to chromosomal origin distribution. In particular, the sequence-specific binding behavior of yeast ORC and this organism’s small genome have allowed genome-wide studies to map the ORC and MCM binding positions and initiation sites of approximately 400 individual yeast origins at near-nucleotide resolution (12; 13; 14; 15; 16; 17). These and other studies have also identified origin-adjacent chromatin features (e.g. nucleosome positioning, modification states, non-histone chromatin-associated proteins) and functional properties (e.g. origin replication time in S-phase, origin efficiency, ORC binding mechanisms) (18; 19; 20; 21; 22; 23). Thus, yeast origins can be parsed by selected criteria into large cohorts whose comparisons can illuminate issues relevant to genome replication dynamics that would be difficult to glean from classical molecular approaches. Such studies have established that approximately half of yeast origins, distributed over the central regions of chromosomes, act in the first portion of S-phase (early and mid origins), while the remaining origins, located near chromosome ends, act later in S-phase (late origins). This origin distribution creates a highly reproducible spatiotemporal pattern of chromosome duplication that can be quantitatively assessed in a population of proliferating yeast (**Figure 1**). Recent reports have defined specific chromatin-associated proteins enriched within the central regions of yeast chromosomes that recruit a limiting S-phase kinase required for MCM complex activation, thus promoting the early activation of origins located within these regions (11; 8). In contrast, it is unclear whether and how chromatin structure impinges on the chromosomal distribution of loaded MCM complexes in yeast.

**Figure 1:**
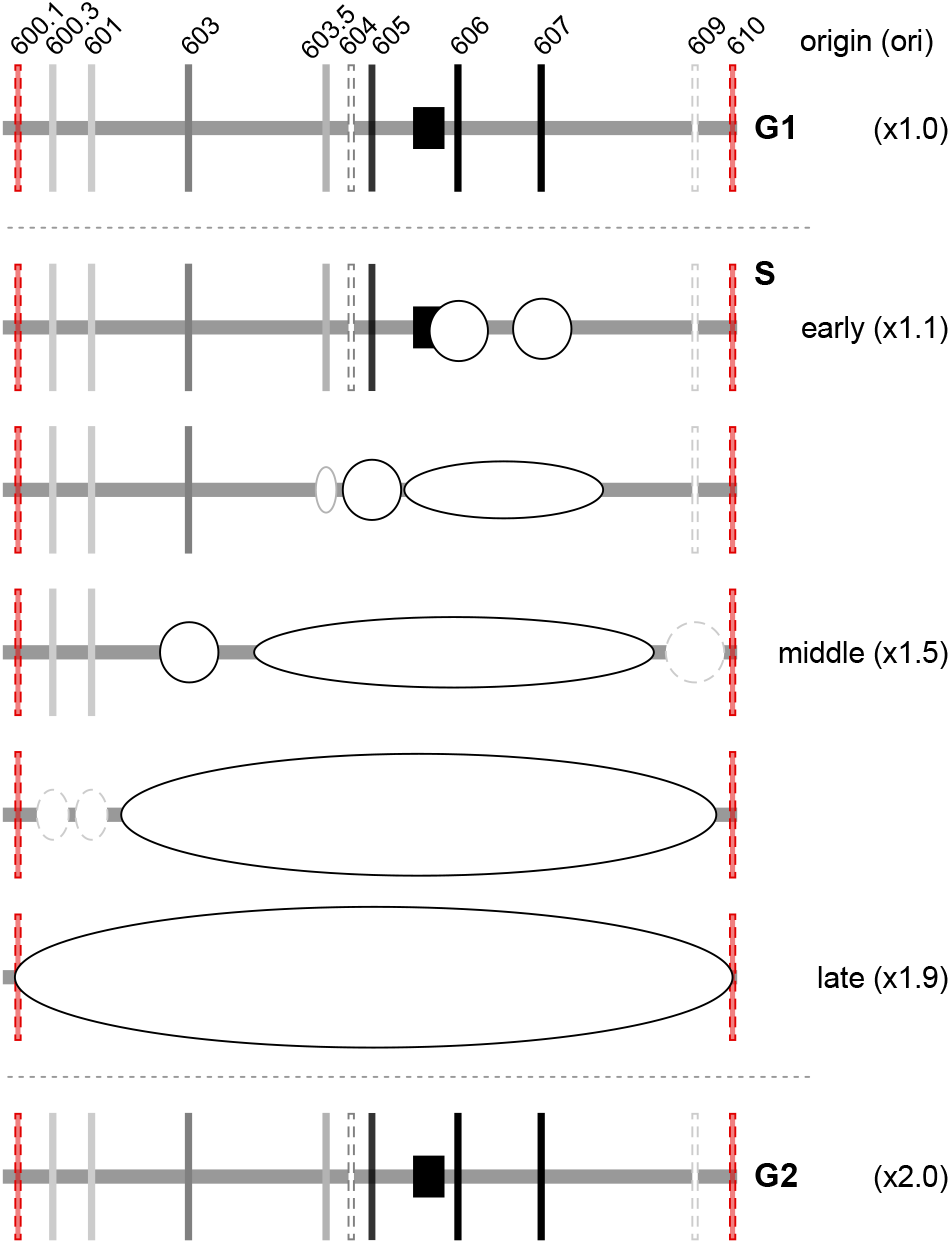
Spatiotemporal progression of yeast chromosome VI replication. Replication of yeast chromosome VI (grey horizontal line) in a population of dividing yeast. Mapped origins are indicated with perpendicular lines. The square represents the centromere. Origins are activated during S phase. The origins with the highest probability of acting are depicted by black vertical lines. These origins are efficient, meaning that they act in most cell cycles. Less efficient origins are depicted in gray, and dotted lines depict the least efficient origins. An inefficient origin is used in only a small fraction of cell cycles. The origins that exist within telomeres are colored red. Telomeres are packaged into Sir2-heterochromatin that represses origin activity (29).

The yeast heterochromatin deacetylase, Sir2, and the heterochromatic nucleosome binding protein Sir3, were recently shown to act directly on nucleosomes adjacent to euchromatic origins, an unanticipated result given the paradigm of Sir proteins as stable components of repressive chromatin that inhibits both transcription and origin function (24; 25; 26). Several independent genetic experiments provide evidence that the Sir2-chromatin molecular signatures found near euchromatic origins are functionally relevant (27; 28; 24). In particular, a deletion of *SIR2* (*sir2*Δ) suppresses the temperature-sensitive growth and MCM complex loading defects caused by the *cdc6-4* mutation. However, because the functional relevance of Sir2 in origin licensing has only been assessed in mutant yeast harboring defective MCM complex loading machinery, the physiological role of Sir2-chromatin in origin licensing is unclear.

We used genome-wide mapping of MCM binding to show that Sir2 promotes a more equitable distribution of origin licensing between early- and late-euchromatic and telomeric-X origins. In *SIR2* cells, these three distinct origin cohorts showed similar levels of MCM binding, whereas in *sir2*Δ cells, telomeric-X and early-euchromatic origins gained MCM relative to late-euchromatic origins. Telomeric-X origins exist within Sir2 heterochromatin, and Sir2-chromatin marks were higher at early- compared to late-euchromatic origins. Thus, Sir2-mediated attenuation of MCM binding at origins correlated with the levels of Sir2-chromatin marks at these elements. Direct assays of replication showed that the enhanced levels of MCM observed at early-euchromatic and telomeric-X origins in *sir2*Δ cells correlated with enhanced replication activity of these elements, providing evidence that their origin activity was limited at the level of MCM complex loading. Finally, in the absence of Sir2, several regions of late-replicating euchromatin failed to complete duplication by the end of S-phase by a mechanism that was independent of the replication capacity of the Sir2-controlled rDNA locus that can indirectly alter euchromatic replication (30; 31). Thus, varying degrees of direct Sir2-mediated attenuation of MCM complex loading at individual origins balances the distribution of licensed origins across chromosomes to ensure the complete duplication of euchromatin by the end of S-phase. These findings describe the first example of a chromatin-mediated mechanism that promotes the spatial distribution of licensed origins across eukaryotic chromosomes.

## Results

### Early origins were enriched among euchromatic origins most responsive to *SIR2*

MCM complex loading at origins requires Cdc6. Therefore, MCM ChIP-Seq signals (MCM signals) are abolished at chromosomal origins in *cdc6-4* cells cultured at 37°C, but restored at many origins, including euchromatic origins, in *cdc6-4 sir2*Δ cells (24). However, not all euchromatic origins in *cdc6-4 sir2*Δ cells are rescued for MCM signals to the same degree. Because Sir2-heterochromatic regions are late replicating and inhibitory to origin function, we initially predicted that the euchromatic origins most affected by *SIR2* (i.e. defined as most effectively rescued for MCM binding in *cdc6-4 sir2*Δ cells compared to *cdc6-4* cells) would be late origins (29; 30; 32; 33). However, instead we found that the euchromatic origins most responsive to *SIR2* were enriched for early origins and depleted for late origins (**Figure** 2**A**).

**Figure 2:**
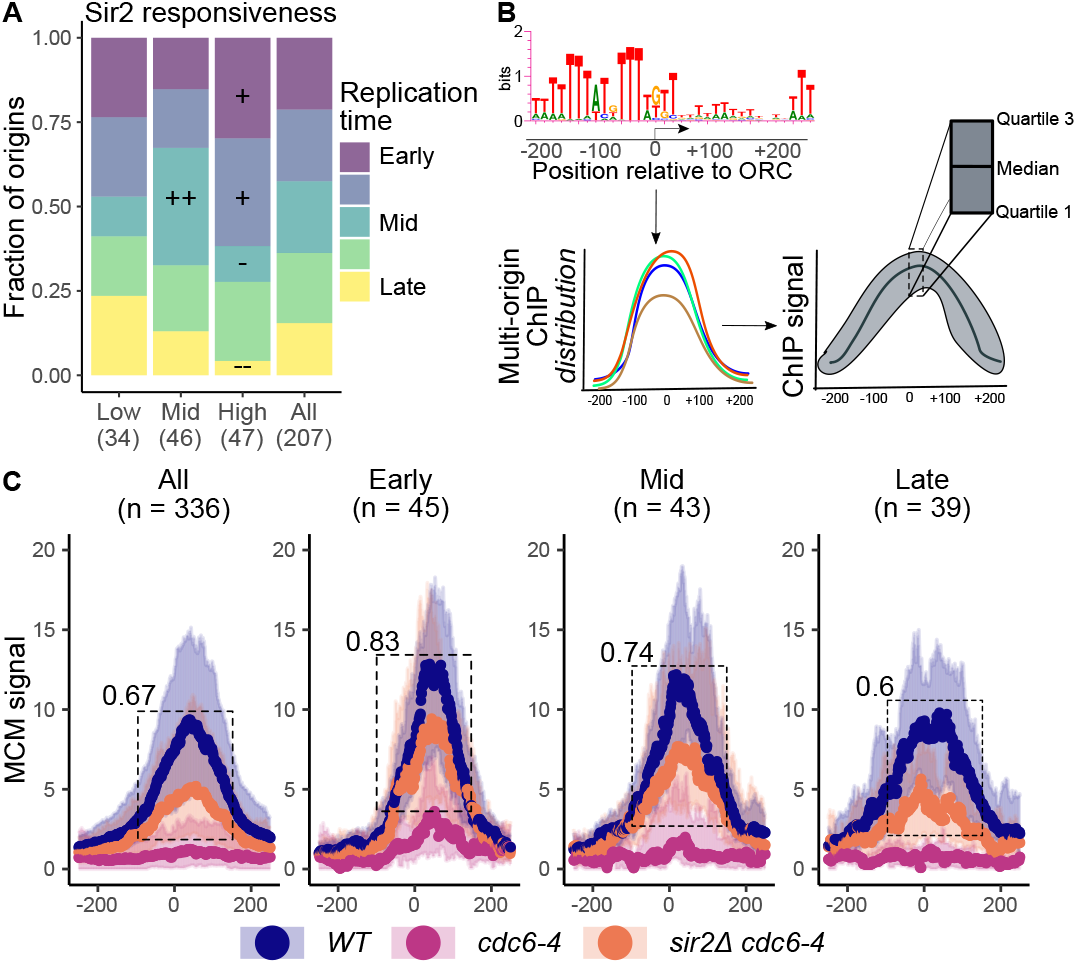
Early origins were enriched among euchromatic origins most responsive to Sir2. **A**. The relationship between Sir2-responsiveness and an origin’s replication time was determined (21). An origin’s Sir2-responsiveness is the ratio of its MCM signal in *cdc6-4* sir2L1 compared to *sir2*Δ cells. Sir2-responsive origins were parsed into quintiles, with the first (Low), third (Mid) and fifth (High) quintiles producing median Sir2-reponsive values of 0.4, 0.7, and 1.0, respectively (24). Euchromatic origins that generated a Trep value in (21) were parsed into quintiles from earliest replicated in S-phase (Early) to latest (Late). Hypergeometric distribution was used to compare the enrichments (+) or depletions (−) within each Sir2-responsive group relative to ‘All’ origins. The P-values are indicated (+/− P<0.05; ++/− P<0.01). **B**. Determining nucleotide-resolution multi-origin distribution of MCM signals generated within origin cohorts (**Figure S1**). The MCM signal associated with a given nucleotide was the median value derived from three biological replicates. To derive the the MCM distribution for the origins within a given cohort, the per nucleotide quartile distribution of MCM signal for all origins in that cohort was determined, plotting the median (dots) and flanking quartiles (shaded coloring). **C**. MCM-signal distributions for wild type (*CDC6 SIR2*), *cdc6-4*, and *cdc6-4 sir2*Δ were determined for each of the indicated Trep groups. The number at the left corner of the box (dotted lines) is the ratio for median MCM signals in *cdc6-4 sir2*Δ to that in *CDC6 SIR2* at each nucleotide summed between −100 through +150.

The analyses in **Figure** 2**A** used MCM signals derived from combining data from three biological replicates and smoothing the data by binning signals from individual nucleotides (24). To examine the data at nucleotide-resolution, we used the approach depicted in **Figure 2B**. MCM signals, while abolished in *cdc6-4* cells, were rescued at euchromatic origins in *cdc6-4 sir2*Δ cells, as expected (**Figure 2C**, All). However, early origins were rescued more efficiently compared to late origins, consistent with the outcome in **Figure 2A**. Thus, while all euchromatic-origin cohorts included Sir2-responsive origins, the early cohort was more likely than the late cohort to contain origins that showed the highest levels of Sir2-responsiveness.

### Sir2-chromatin marks were higher at early compared to late euchromatic origins

Molecular hallmarks of Sir2 heterochromatin, Sir2-dependent depletion of H4K16ac and Sir3 binding to nucleosomes, are present at euchromatic origins (24). Therefore, we asked whether early- and late-euchromatic origin cohorts showed differences in the levels of these molecular signatures (**Figure** 3). While both early- and late-euchromatic origin cohorts were flanked by nucleosomes showing Sir2-dependent depletion of H4K16ac as well as Sir3 binding, the early-origin cohort showed greater levels of these marks compared to the late-origin cohort. The difference was most pronounced at the −1 and +1 nucleosomes flanking the ORC site. Thus the origin cohorts’ levels of Sir2-chromatin marks correlated with their Sir2-reponsiveness.

**Figure 3:**
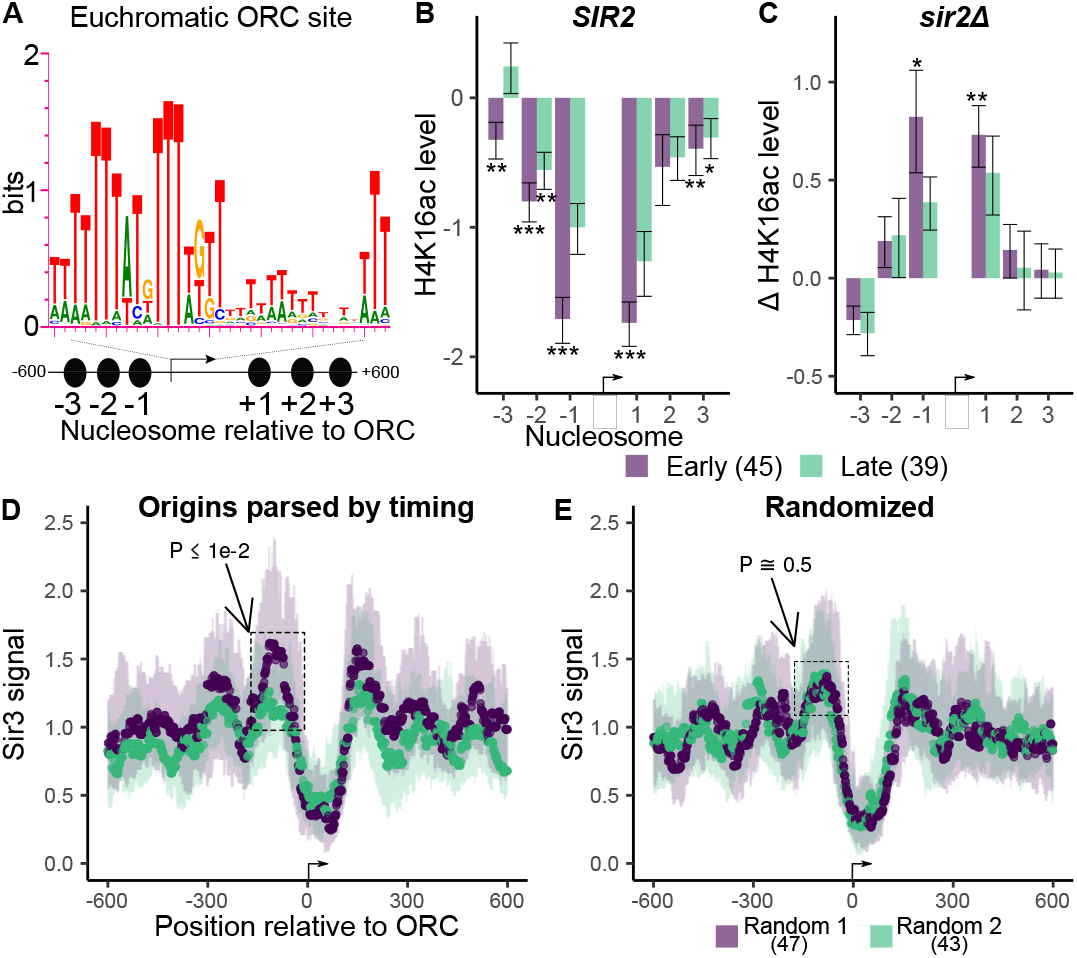
Sir2 chromatin marks were higher at early compared to late euchromatic origins. **A**. Early- and late-euchromatic origin cohorts were examined for Sir2-chromatin marks on the six origin-adjacent nucleosomes, as indicated (24). **B**. Normalized H4K16ac assessed in *SIR2* cells at early- and late-euchromatic origins. **C**. The change in H4K16ac observed for early- and late-euchromatic origin-adjacent nucleosomes in *sir2*Δ cells. **D**. Sir3 MNase ChIP-Seq signals at early- (purple) and late-euchromatic (mint) origin cohorts. Boxed is a nucleosome upstream of the ORC site where the median Sir3 signal at early origins differed significantly (Wilcoxon rank sum tests) from that at late origins. The significance cut-off demanded that > 90% of positions within signal apexes had P-values < 0.05. The region marked had a mean P-value ≤ 0.01. 0.01. **E**. The origins in the early- and late-origin cohorts were randomly assigned to two new groups and the Sir3 MNase ChIP-Seq signals from (34) were determined for each new randomized cohort. The same region of significant divergence in Sir3 signals in D is indicated as well as the mean P-value.

### *SIR2* promoted more equitable distribution of MCM complexes between early and late euchromatic origins

*SIR2* has a profound effect on MCM complex loading at euchromatic origins in *cdc6-4* cells, but its effect in wild-type (*CDC6*) cells was less clear (**Figure 2C**) (24). Therefore, we used the approach described in **Figure 2B** to compare MCM signals between different origin cohorts in *CDC6* cells that differed in their *SIR2* genotype. (**Figure 4**). In *SIR2* cells, the late-euchromatic origin cohort generated lower median MCM signals than the early-euchromatic origin cohort, consistent with a previous study (10). However, a substantial overlap in the distribution of signals for these two cohorts was observed, suggesting that the MCM binding differences between early- and late-euchromatic origin cohorts were minimal (**Figure 4A**). In contrast, in *sir2*Δ cells, the difference in MCM signals between these same two cohorts was enhanced and significant (see also **Figure 4C**). Indeed, MCM signals at >50% of the nucleotides spanning (−10 to +100) differed significantly between late- and early-origin cohorts in *sir2*Δ cells, whereas no single nucleotide signal differed significantly in *SIR2* cells (P<0.01) **(Figure S2)**. MCM signals were also assessed at telomeric-X origins, elements known to be repressed by Sir2-dependent heterochromatin (29). In *SIR2* cells, telomeric-X origins generated MCM signals similar to the euchromatic-origin cohorts, whereas in *sir2*Δ cells, the telomeric-X origin cohort generated MCM signals that were more similar to those of the early-origin cohort and greater than the late-origin cohort (**Figure 4B**). Therefore, Sir2 limited MCM levels at both telomeric-X origins packaged into Sir2-dependent heteorochromatin as well as early-euchromatic origins relative to late-euchromatic origins.

**Figure 4:**
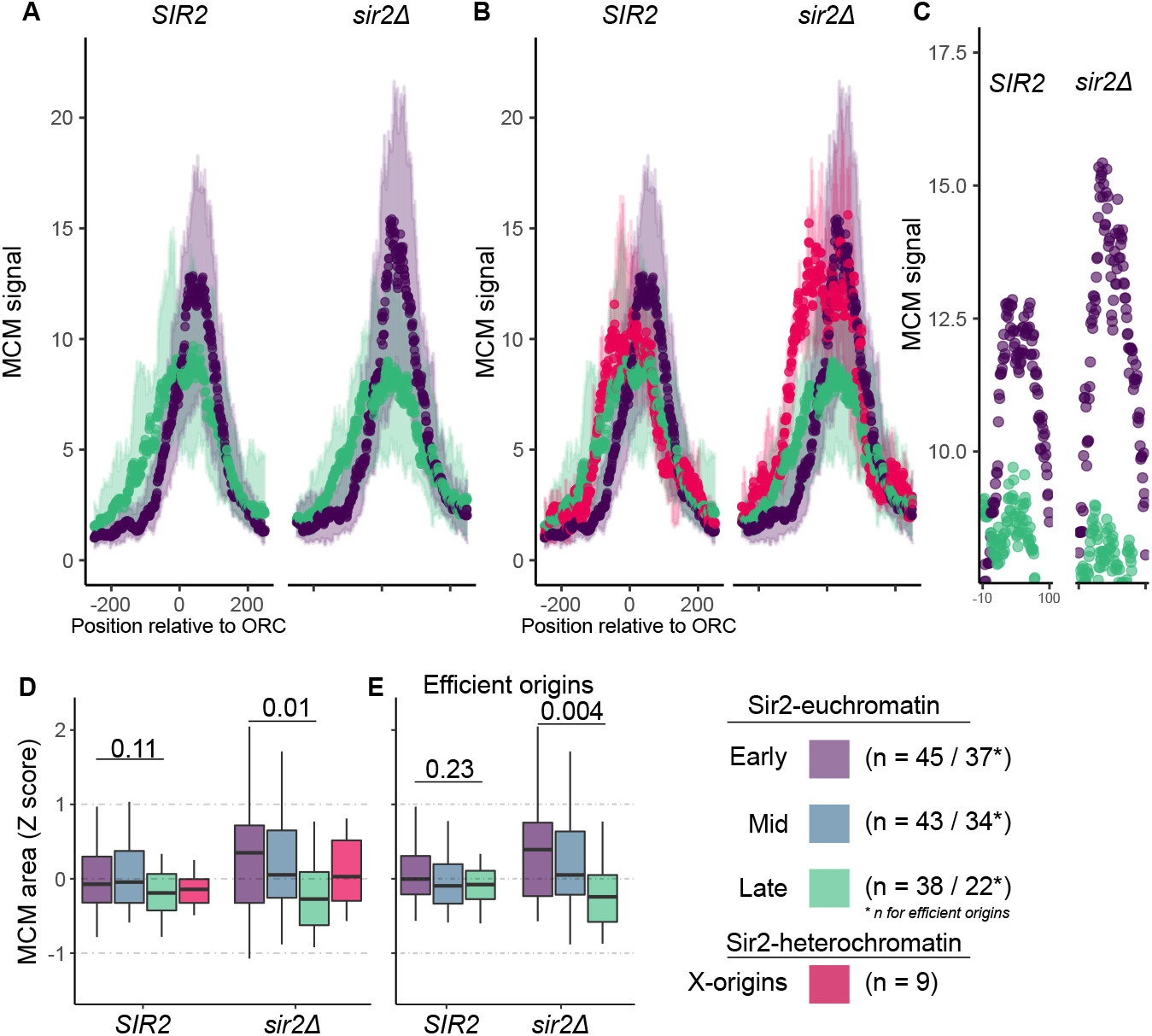
*SIR2* promoted more equitable distribution of MCM complexes between early- and late-euchromatic origins. **A**. MCM signals were determined as in **Figure** 2**C** for the indicated origins in congenic *SIR2 CDC6* and *sir2*Δ *CDC6* cells. **B**. Same analyses as in A, except data for the nine telomeric-X origins also shown. **C**. Magnified view of the median values between nucleotides −10 and +100 for the early- and late-euchromatic origin cohorts in A. **D**. Median signals between nucleotides −10 and +100 were summed to generate an MCM area for each origin. Z-scores were then assigned to each origin and displayed in box-and-whiskers plots for each indicated cohort. The P-values (Wilcoxon rank sum test) for the difference between the early- and late-origin cohorts’ median MCM Z-scores are indicated. **E**. Analysis as in D but with exclusion of inefficient origins (17).

To more easily quantify differences between origin cohorts, the MCM signals between the −10 and +100 nucleotides for each of the fragments were summed, and the sums used to assign each origin a Z-score to indicate how far the MCM signal for that origin diverged from the mean behavior of the entire collection of origin fragments (the mean is defined as ‘0’). The Z-scores for each cohort were displayed in box-and-whiskers plots (**Figure 4D**). In *SIR2* cells, each of the origin cohorts generated similar MCM signals, indicating that MCM complexes were equitably distributed among these cohorts. In contrast, in *sir2*Δ cells, the median Z-score of the early-origin cohort rose above the mean score in the population, while the median Z-score for the late-origin cohort fell below the mean, such that the difference between the MCM signals generated by the early- and late-euchromatic origin cohorts was clear and significant. The telomeric-X origin cohort also generated higher MCM Z-scores in *sir2*Δ cells compared to the late-euchromatic origin cohort. Thus, in the absence of Sir2, MCM complexes were more efficiently distributed to early- and mid-euchromatic and telomeric-X origins compared to late euchromatic origins. The Sir2-controlled rDNA origin did not show enhanced MCM association in *sir2*Δ cells, suggesting that MCM complex loading efficiency of the rDNA origin is not affected by Sir2, as reported previously **(Figure S3**) (35).

Some of the origins within each of these cohorts act in only a small fraction of cell cycles, possibly because they are licensed inefficiently (17). To focus on only those origins that were most likely to be licensed, we repeated the analysis using only those origins within each cohort that generated a measurable origin-efficiency value in a genome-scale analyses of origin function (17). Considering only these origins, the difference between MCM levels at the early- and late-origin cohorts observed in *sir2*Δ cells was enhanced. (**Figure 4E**). Thus, Sir2 prevents MCM complexes from accumulating within early-relative to late-replicating euchromatin.

### Changes in MCM distribution correlated with changes in replication dynamics

To address whether the alterations in MCM distribution described above had the potential to alter replication dynamics, S-phase Sort-Seq experiments were performed. These experiments used the same yeast strains as for MCM ChIP-Seq in **Figure** 2, except that cells were cultured at the permissive growth temperature for *cdc6-4* so that the effect of this mutation itself on replication dynamics could be examined. In S-phase Sort-Seq, the number of sequence reads for a given region in S-phase are normalized to the corresponding reads from G1-phase to generate a Sort-Seq value (36). Sort-Seq values annotated to a chromosome produce a replication profile where peaks indicate active origins (**Figure S4**). Wild-type and *sir2*Δ cells generated virtually indistinguishable profiles, indicating that any Sir2-mediated effects on euchromatin replication dynamics were not observable at this level of resolution. In contrast, *cdc6-4* cells, regardless of *SIR2* genotype, showed reduced origin function across several regions of the genome. Thus, only a fraction of chromosomal origins functioned normally in *cdc6-4* cells even when these cells were cultured at permissive temperature, and the *SIR2* genotype did not affect origin preference. Notably, Sir2 did not alter the origins that remained the most functional when MCM complex loading was compromised by *cdc6-4*.

To enhance comparison of the relevant origin cohorts, Sort-Seq values were assigned to 30 kb replicons, the replicons parsed into cohorts by their origins’ replication times (Treps) or Sir2-responsiveness, and the data presented in box-and-whiskers plots (**Figure 5A-B**). The results for *CDC6* cells validated the effectiveness of this approach: the early-replicon cohort generated greater Sort-Seq values than the late-replicon cohort, and Sir2 delayed replication of the telomeric-X origin cohort, as expected. In *cdc6-4* cells, regardless of *SIR2* genotype, the relative behavior of these replicons was maintained, though the variation between cohorts and within the late-origin cohort was compressed. However, when these replicons were parsed based on Sir2-responsiveness, a *cdc6-4* effect on origin usage was observed (**Figure 5B**). First, in *CDC6* cells, regardless of *SIR2* genotype, the three distinct Sir2-responsive cohorts produced similar broad box-and-whiskers plots, an expected outcome given that each of these cohorts contained a mixture of replicons duplicated at different times in S-phase (**Figure 2A**). However, in *cdc6-4* cells, regardless of *SIR2* genotype, the different Sir2-responsive cohorts produced box-and-whisker plots similar to those produced for the cohorts parsed by their S-phase replication times (compare *cdc6-4* data in **Figure 5A-B**). Thus, in *cdc6-4* cells, the most Sir2-responsive origins showed the highest median replication activity in S-phase. Sir2-reponsiveness is defined by the level of MCM complex loading *cdc6-4 sir2*Δ cells. Therefore, the origins most effective at binding MCM when origin licensing was compromised by the *cdc6-4* mutation were the same origins that showed the highest replication function in *cdc6-4* cells, providing evidence that alterations in MCM distribution had an impact on replication dynamics.

**Figure 5:**
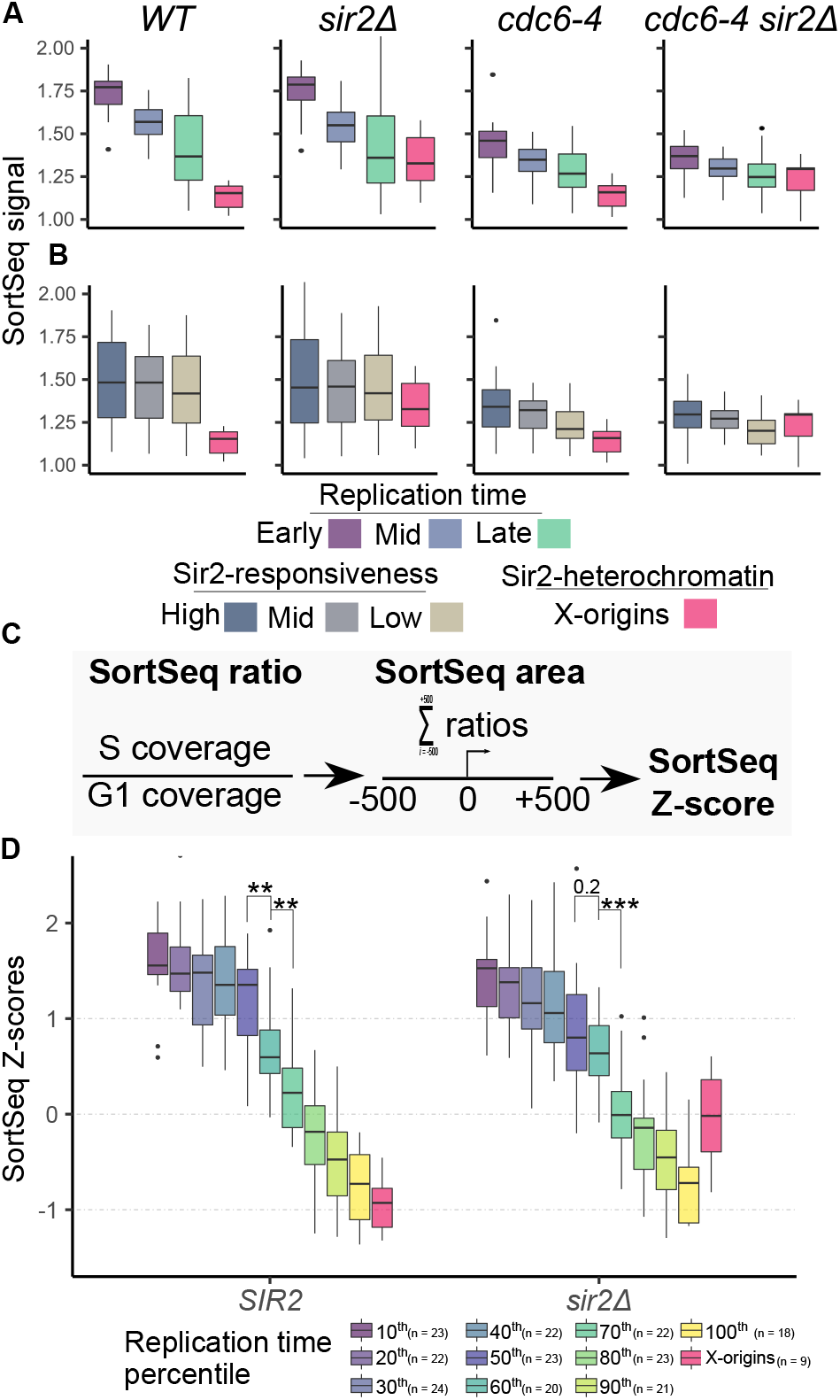
Changes in MCM distribution coincided with corresponding changes in replication dynamics. **A**. Sort-Seq values for 30 kb replicons centered on origins parsed by their Trep values from (21). Telomeric-X replicons also shown. **B**. As in A except with replicons parsed by euchromatic origins’ assigned Sir2-responsiveness. (24). **C**. Assigning Sort-Seq Z-scores to distinct 1001 bp origin fragments. **D**. Euchromatic origins parsed into ten distinct cohorts by their Trep values. Z-scores are plotted from the earliest 10% (10^th^) to the latest 10% (100^th^) decile in *SIR2* and *sir2*Δ cells. Telomeric-X origin Z-scores are also plotted. The Wilcoxon rank sum P-values for the differences between the 50th and 60th and 70th cohorts are indicated with **< 0.01 and ***<0.001.

### Sir2 promoted the normal progression of euchromatic origin replication in an unperturbed S-phase

The above analyses suggested that in *CDC6* cells, Sir2 did not affect euchromatic origin function in S-phase, but the dynamic range of the analyses was limited in part by DNA replication itself (1n-2n). In the spatiotemporal control of chromosome duplication, a more meaningful parameter is how specific genomic regions are duplicated relative to one another (37). Therefore, to examine the replication behavior of origin cohorts at higher-resolution and over a greater dynamic range, we assigned Z-scores to defined origin fragments based on their calculated Sort-Seq ratios (**Figure** 5**C**), ranked the origins by their S-phase replication time, portioned the ranked origins into deciles and displayed the Z-scores for each decile in box-and-whiskers plots (**Figure** 5**D**). This approach revealed effects of Sir2 on euchromatic replication that were not apparent from the previous analyses. First, in *SIR2* cells, the 50th and 60th deciles (early/mid-S origins) differed significantly, whereas they did not in *sir2*Δ cells. Second, the median values for the early-origin deciles (10th-50th) in *SIR2* cells were similar, whereas they continually decreased in *sir2*Δ cells, indicating that the replicative competitiveness of individual origins within these cohorts was diverging, with the earliest cohorts gaining in relative replication efficiency (**Figure S5**). Third, the difference in the replication behavior between the 60th and 70th ranked cohorts was enhanced in *sir2*Δ cells, creating a clear gap in their ranked replication function. Thus, *SIR2* promoted the normal progression of euchromatic origin function in an unperturbed S-phase.

### *SIR2* was required for completing replication of late replicating euchromatin independent of rDNA replication demands

Analyses of our S-phase Sort-Seq data revealed that Sir2 had a detectable effect on the replication progression of euchromatic origins, with later-euchromatic origins falling further behind both early-euchromatic origins and telomeric-X origins in *sir2*Δ cells (**Figure** 5**D**). These effects might be relevant to the inability to complete the duplication of late-replicating euchromatin by early-G2 phase in *sir2*Δ cells as documented in (31). Therefore, the euchromatic- and telomeric-X origin cohorts were assessed using the data sets from this previous study. When the Z-score approach was applied to the S-phase Sort-Seq data from this independent study, similar alterations in the relative behaviors of the 50th, 60th and 70th cohorts were observed, strengthening the conclusion that the phenomenon was *SIR2*-dependent (**Figure S6**). Next, the effect of Sir2 on the replication of Trep-parsed euchromatic origin cohorts was examined (**Figure** 6**A**). These origin cohorts produced the expected box-and-whiskers plots in S-phase in both *SIR2* and *sir2*Δ cells. However, in the G2-phase populations, the *SIR2* and *sir2*Δ cells generated different outcomes. In *SIR2* cells, early- and late-euchromatic origin cohorts generated indistinguishable Sort-Seq Z-scores near ‘0’, indicating that these fragments had been equivalently duplicated. In contrast, in *sir2*Δ cells the behavior of these two origin cohorts diverged in the G2-phase population, with the majority of early origin-fragments generating Z-scores above the mean of the population, and the majority of late origin-fragments generating Z-scores below the mean (**Figure** 6**A**). Thus, while in *SIR2* cells, only the telomeric-X origins remained unduplicated in G2-phase, in *sir2*Δ cells both late-euchromatic origins and telomeric-X origins remained unduplicated. Thus Sir2 was required to complete the duplication of late-euchromatic origin fragments by early G2-phase.

**Figure 6:**
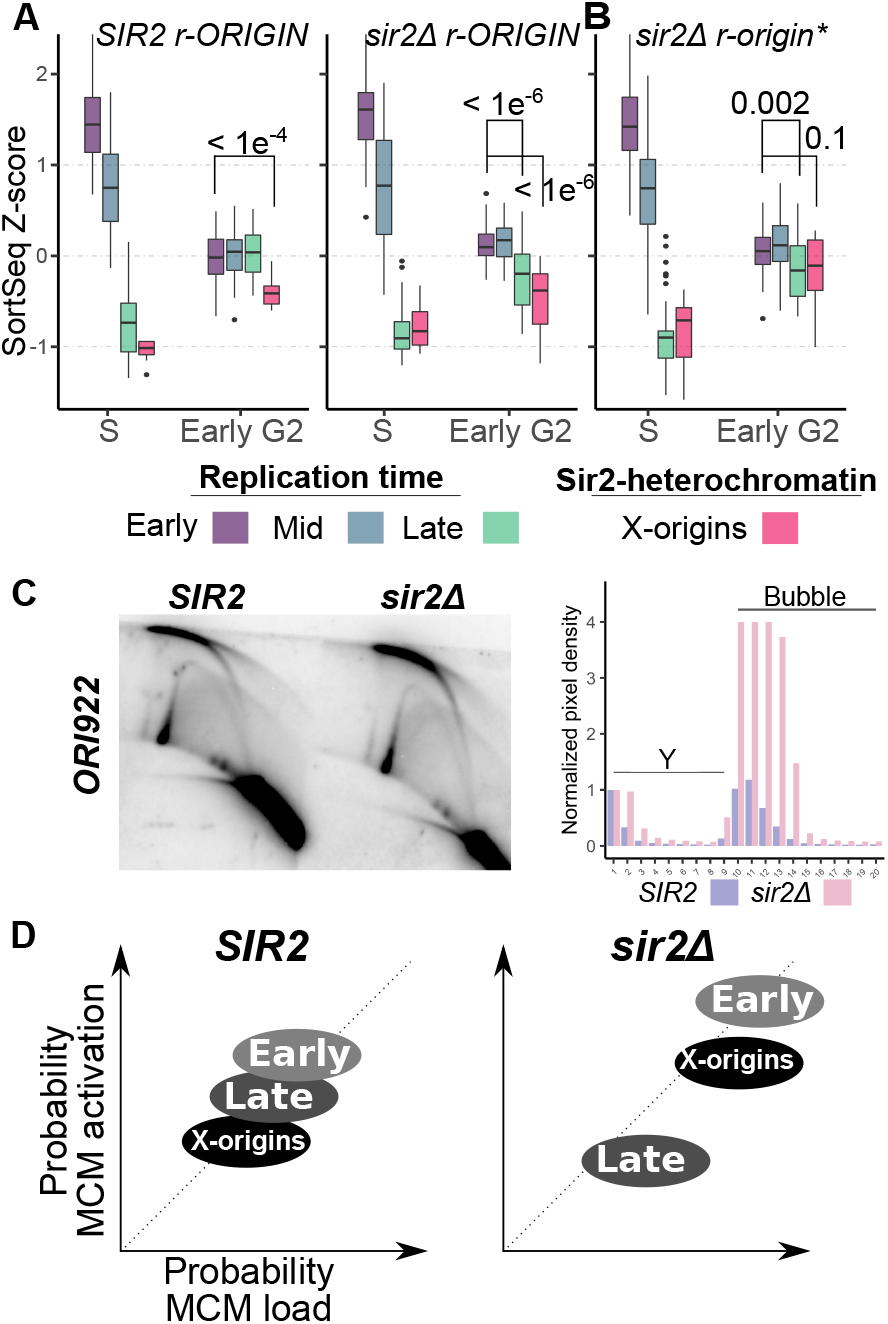
*SIR2* was required for completing replication of late euchromatic regions independent of rDNA replication demands. **A**. Z-scores for the indicated origins from S-phase and early G2-phase *SIR2* and *sir2*Δ cells derived from raw data in (35) were determined as in **Figure 5C** and displayed as box-and-whiskers plots for early-, mid-, and late-euchromatic origins, and telomeric-X origins. **B**. Analyses as in 6A except with data from *sir2*Δ *r-origin\** mutant. **C**. Two-dimensional origin mapping of *ORI922* in indicated strains (left panel). Quantitation of these data (right panel). Radioactive intensities for each indicated region were normalized by dividing signals by that generated by the largest replication forks (**Figure S7**). **D**. Sir2 controls the relative replication probability of the three indicated origin cohorts at the level of origin licensing efficiency (probability of MCM complex loading) and S-phase origin activation (probability of MCM complex activation). While these two steps are distinct, because the MCM complex is the substrate for the S-phase origin activation factors, alterations in MCM complex loading efficiencies in turn affect the ability of the cohorts to compete for limiting S-phase origin activation factors.

Sir2 can indirectly affect early euchromatic origins via its direct repression of the rDNA origin that is present in hundreds of copies within the yeast rDNA repeat array (30). Specifically, in *sir2*Δ cells, the rDNA origins (r-ORIGIN) are more active, and because there are so many, limiting S-phase MCM complex activation factors get sequestered away from euchromatic origins causing a delay in euchromatic-origin activation. This *sir2*Δ-caused sequestration of S-phase MCM complex activation factors contributes to the incomplete duplication of late replicating regions in G2 phase (31). In support of this model, *sir2*Δ cells harboring a mutation within the rDNA origin, referred to here as *r-origin\**, reduces rDNA-dependent sequestration of S-phase MCM complex activation factors and partially rescues the delayed replication of late regions caused by *sir2*Δ. If the changes in MCM complex distribution described here were relevant to the incomplete duplication of late-euchromatic origins, then the *sir2*Δ-caused under replication of these euchromatic elements should not be rescued by the *r-origin\** mutation. In contrast, the delayed duplication of telomeric-X origins that had gained MCM complexes in *sir2*Δ cells should be (see **Figure** 4). Therefore the replication of the relevant origin cohorts was assessed in *sir2*Δ *r-origin\** cells (**Figure 6B**). In *sir2*Δ *r-origin\** cells, the duplication of the telomeric-X origin cohort in early G2 was substantially enhanced, such that the difference between early-euchromatic origins and telomeric-X-origins in early G2 no longer reached a significant P-value cut-off (**Figure 6A-B**, compare early euchromatic and telomeric-X origins in early G2 in *sir2*Δ *r-ORIGIN* cells to their relative behavior in *sir2*Δ *r-origin\** cells). Therefore, the duplication of telomeric-X-origins in *sir2*Δ cells was limited by the availability of S-phase MCM complex activation proteins sequestered by the rDNA array (31). However, in contrast, the late-euchromatic origin cohort remained under duplicated even in the *sir2*Δ *r-origin\** cells (**Figure 6A-B**, in particular compare *sir2*Δ *r-origin\** to *SIR2 r-ORIGIN*). Specifically, more than half of the late-euchromatic origins remained incompletely duplicated in early G2-phase despite increased availability of of S-phase MCM complex activation factors. These data provided evidence that the reduced licensing of late-euchromatic origins relative to other origin cohorts contributed to their *sir2*Δ-dependent delay in duplication. Thus, Sir2-enforced distribution of MCM complexes across chromosomes insured that late euchromatin was duplicated by the end of S-phase.

### Sir2 limited both MCM binding and origin function at an early, efficient euchromatic origin, *ORI922*

The analyses the MCM ChIP-Seq data indicated that *SIR2* prevented early-euchromatic origins from gaining MCMs relative to late-euchromatic origins. If the relative gains in MCM signals in *sir2*Δ cells were indeed due to more cells in the population having effectively licensed these euchromatic origins, then the function of these origins should be enhanced in *sir2*Δ cells. The quantitative analyses of Sort-Seq data indicated that many early- and mid-euchromatic origins generated higher ranking Z-scores in *sir2*Δ compared to *SIR2* cells, consistent with this intepretation. To address this issue using an independent method, two-dimensional origin mapping was used to examine the chromosomal euchromatic origin *ORI922 (38)* (**Figure 6C** and **Figure S7**). Two-dimensional origin mapping allows a direct assessment of initiation efficiency by examining the levels of bubble intermediates relative to replication forks. *ORI922* is an early and relatively efficient origin (Origin Efficiency Metric = 0.75 (17). An OEM of 1.0 is assigned to the most efficient origin in the genome, *ORI305*). We reasoned that *ORI922* had enough capacity to improve in function to allow detection of enhanced efficiency by origin mapping. As expected, *ORI922*, was an efficient origin in *SIR2* cells. Nevertheless, in *sir2*Δ cells, an enhancement in origin activity of *ORI922* could be detected (**Figure 6C**). Therefore, Sir2 chromatin at *ORI922* limited its origin activity at the level of MCM complex loading (**Figure S8**)

## Discussion

Accumulating evidence indicates that the yeast Sir proteins, non-essential chromatin regulators best known for their roles in forming yeast heterochromatin (a.k.a. transcriptionally silent chromatin), also act directly within euchromatin where they can have an impact on chromosome replication (39; 26; 24). In particular, Sir2 and Sir3 act directly on the 2-4 nucleosomes immediately adjacent to euchromatic DNA replication origins where they can inhibit MCM complex loading (24). However, the functional potential of this Sir2-chromatin state to inhibit MCM complex loading has thus far only been assessed in cells in which one of the core origin licensing proteins is defective (e.g. *cdc6-4*, *orc5-1*, *mcm2-1* mutant yeast) (27; 28; 24). Thus, a physiological role for Sir2 chromatin on MCM complex loading in wild-type yeast was unclear.

In this study, genomic and computational approaches documented a clear and statistically significant shift in MCM complexes toward early-replicated euchromatin in the absence of Sir2. The higher levels of Sir2-chromatin marks at early- compared to late-euchromatic origins explained why the Sir2-attenuating effect on MCM complex loading was on average more robust at early- compared to late-euchromatic origins. Sir2’s inhibitory effects on MCM complex loading depends on its catalytic deacetylation of nucleosomes (27; 28; 24). Therefore, we propose that the repressive properties of Sir2-chromatin (e.g. reduced DNA accessibility, reduced nucleosome mobility) act to attenuate MCM complex loading by reducing the probability that an origin licensing reaction will be completed. The probability of any one or several of a number of discrete steps–e.g. ORC binding, Cdc6 binding, Cdt1-MCM association etc–could be affected, and to varying degrees at individual origins. The higher levels of Sir2 chromatin at Sir2-responsive origins indicate that the levels of this repressive chromatin-state are a component that affects the probability of origin-licensing, but other Sir2-independent variables such as ORC-DNA affinity or nucleosome orientation could also be relevant, establishing a level of stochasticity to this attenuating mechanism (24). In terms of cell physiology, the key point is that relatively small, stochastic reductions in the probability of origin licensing at individual origins accumulates over multiple origins to impose a significant genome-scale impact: origin density is concentrated in early-replicating euchromatin and at telomeric-X origins and thereby relatively depleted from late-replicating euchromatin. Essentially, Sir2 prevents early-replicating euchromatin from being ‘too effective’ at origin licensing, thus limiting the colllective origin activity that would otherwise be possible in these regions (**Figure 6D**). Indeed, because MCM complexes are the substrate for the limiting S-phase origin activation factors, and early-replicating euchromatin has active mechanisms for recruiting these factors, a shift in MCM complexes to early-replicating euchromatin will lead to a concomitant shift in S-phase origin activation factors. The result is more origin activity in early-replicating euchromatin than required for its efficient duplication, and a depletion of origin activity from late-replicating euchromatin, increasing the probability that late-replicating euchromatin will not complete duplication within S-phase (**Figure** 6**D**).

While Sir2-heterochromatin-mediated inhibition of origin function at telomeres is an established phenomenon, it was not known whether this inhibition occurs at the level of MCM complex loading or S-phase activation (29). Thus, an important observation was that Sir2 attenuated MCM complex loading at heterochromatic telomeric-X origins substantially, although even at these heterochromatic domains, MCM complex loading was not completely prevented by Sir2. Regardless, the increased probability of licensing telomeric-X origins in *sir2*Δ cells provided an explanation for how these regions competed more effectively than late-euchromatic origins for S-phase MCM complex activation factors (**Figure 6D**). Thus, in the absence of Sir2, both early euchromatic and telomeric X-origins were more competitive for MCM complexes than late-euchromatic origins, and in turn were more competitive for the limiting S-phase origin activation factors. Notably, the rDNA origin did not show enhanced levels of MCM in *sir2*Δ cells, consistent with an earlier report, indicating a distinct mechanism for Sir2-mediated inhibition of rDNA origin function at this locus (35).

Early origins exist within the most ‘open’ and transcriptionally active regions of the genome that bind proteins that actively recruit origin activation proteins (18; 11; 8; 3). Thus, it was initially perplexing that Sir2-chromatin, known for its ability to inhibit protein-DNA interactions, was present at higher levels and had a greater negative impact on MCM complex loading at early- compared to late-euchromatic origins. A plausible explanation is that the same molecular properties that make early euchromatic origins more accessible to the DNA replication machinery also make these elements more accessible to Sir proteins that are untethered from heterochromatin and moving through the nucleoplasm. While Sir proteins are concentrated within heterochromatin domains, heterochromatin is dynamic, with Sir proteins releasing and rebinding to these regions (40; 26; 41). Obviously, any released Sir proteins will have opportunities to sample other regions of the genome. The greater accessibility of early euchromatin, as well as ORC’s ability to contribute to Sir-nucleation events within heterochromatin, might facilitate the sampling of early-euchromatic origins by these ‘free’ Sir proteins more frequently compared to less accessible late-replicating euchromatic origins (42; 25; 43).

Classic yeast suppressor genetics uncovered Sir2-chromatin’s negative effect on MCM complex loading, a negative regulatory phenomenon that, like other forms of negative regulation that affect DNA replication, would have been extremely difficuletto uncover without the aid of genetics (27; 44). Building on this foundation, the genomic and computational approaches used here exploited the deeply annotated yeast genome to quantitatively analyze large cohorts of precisely mapped yeast origins. The ability to examine origin-cohorts was essential to understanding the physiological relevance of Sir2-mediated attenuation of MCM complex loading at individual origins. In a single-celled eukaryotic microbe like yeast, such a genome-scale effect could impinge on organismal fitness by reducing cell proliferation rates and/or genome stability, as delayed replication is associated with enhanced mutation frequency (45; 46). However, in mammalian cells, SirT1, the human ortholog of yeast Sir2, as well as Set8-mediated chromatin compaction limit origin function and promote genome stability (47; 48; 49). Notably, the Set-8 mechanism works at the level of limiting MCM binding to chromatin. Thus, genome-scale control of origin activity through repressive chromatin is emerging as an important mechanism for controlling eukaryotic chromosome duplication and stability. The high-resolution information about origin positions and functions available in budding yeast allowed us to show how such a mechanism was important for the normal distribution of origins across chromosomes. While the importance of origin distribution to genome stability has become clear, this is the first study to identify a specific chromatin-mediated mechanism that establishes the spatial distribution of MCM complexes across a eukaryotic genome (50; 51; 52).

## Materials and methods

### Data analysis

All data analyses were performed in R. Scripts are available upon request.

### Yeast Strains

The strains used in this study were congenic derivatives of W303-1A, and were described in detail in (24).

### Sequencing data

The raw data used for the experiments in this study have been assigned the following BioProject ID’s: PRJNA428768 and PRJNA601998 contain the reads from the MCM ChIP-Seq experiments; PRJNA601998 contains the reads from the S-phase Sort-Seq experiments (**Figure 5**). The GEO accession GSE90151 contains the reads from the S and G2 Sort-Seq experiments (**Figure 6**).

### Determining per-nucleotide MCM ChIP-Seq signals and distributions

MCM ChIP-Seq experiments were performed on three independent cultures of each indicated strain (except for for *cdc6-4*) as described (24). For each independent experiment, reads were mapped to sacCer3 using Bowtie2 and default parameters. Duplicates were marked and removed using MarkDuplicates within Picard tools. Per-nt coverages were determined from mapped, unique reads using Samtool’s BedCov. Coverages for each ChIP and input experiments were normalized for sequencing depth and breadth as outlined (53). ChIP/input ratios for each independent culture per strain were mapped to the nucleotides within origin-containing fragments oriented relative to the T-rich ORC site (assigned nucleotide position ‘0’) (**Figure S1**). Once mapped, the ChIP/input ratio for each nucleotide within each replicate was scaled by dividing each ratio by the median ChIP/input ratio measured between coordinates −600 to −400 relative to the start of the ORC site as described for **Figure 2**. Therefore, each locus within each replicate was internally scaled. To derive the per nucleotide MCM ChIP-Seq signal distribution for all origins within a given group (e.g., Sir2-responsive group, Replication timing group), we determined the per nucleotide median ± one quartile of the distribution of scaled MCM ChIP/input ratios measured at that nucleotide position. Thus, every nucleotide is represented as a “whisker-less” box-and-whiskers plot for all origins within a group, reflecting the distribution of signal measured within a given cohort.

### Determining H4K16ac levels at nucleosomes adjacent to euchromatic origins

Normalized H4K16ac levels at relevant nucleosomes were determined as described previously (24). The H4K16ac IP/input ratios for each nucleotide within the coordinates of each nucleosome annotated in (54) were summed. With genome-wide nucleosomal H4K16ac levels thus calculated for every nucleosome in the genome, we mapped the nucleosomes 5’ and 3’ to the T-rich ORC binding site (nucleosome −3 through nucleosome +3) to 1.2 kb fragments centered and aligned to the start of the T-rich strand of the fragment’s ORC binding site. Nucleosomes were also mapped to a control group of loci, 391 randomly selected distinct euchromatic 1.2 kb regions that were not annotated in the OriDB and that lacked an ORCACS match. An arbitrary midpoint was assigned to these control loci and nucleosomes −3 to +3 from this midpoint were mapped. For each control fragment, the mean H4K16ac level for the six nucleosomes was determined (spanning signal), and then the mean of the 391 spanning H4K16ac signals was determined. This value was used to normalize the H4K16ac levels for each nucleosome surrounding the experimental loci **Figure 3**.

### S-phase Sort-Seq experiments

The S-phase Sort-Seq experiments in **Figure 5** were performed on the indicated yeast growing at the permissive-growth temperature for *cdc6-4* (23°C) using the methods described in (36).

### Calculation of Sort-Seq Z-scores

Z-scores were calculated from the sum of G1-normalized Sort-Seq ratios (S / G1 or early G2 / G1) within 1 kbp fragments from unique chromosomal loci. The robustness of the Z-score approach for comparing the relative behavior of independent 1 kb origin fragments within the genome depends on using a large number of 1 kb regions both actively and passively replicated within the population. Therefore, the number of relevant 1 kb regions was maximized by including all unique 1 kb regions outlined in **Figure S1**(n = 1015) that contained a match to the ORC consensus site, regardless of whether that site corresponded to a functional origin. Additionally, 525 additional loci lacking ORC consensus sites and not overlapping the 1015 fragments with matches were included (non-matched loci). For these loci a midpoint was randomly assigned for centering the 1 kbp fragment. Fragments that contributed to Z-scores were chosen irrespective to their presence in euchromatin or heterochromatin.

**Two-dimensional origin mapping of *ORI922*** was performed as described (55; 56) with the following alterations: CsCl purified DNA was generated from three liters of yeast growing in liquid YPD at 30°C, and the three pools of gradient-pulled genomic DNA were combined into a single Falcon tube prior to spooling with isopropyl alcohol. This approach was required to recover ̃100 μg genomic DNA per liter of yeast, which was the amount of DNA used for an experiment. Three radioactive probes were generated by three separate random labeling reactions (α^32^P-dCTP) of three distinct gel-purified fragments generated by PCR of yeast genomic DNA. The primer pairs used to generate the *ORI922* probe fragments were: Pair (1) Fwd, TCCTTGGTAGCAAAGATAAAG, Rev, CTGGGAAACCAAAAGATC; Pair (2) Fwd, TAACATGGGTACAGCAAATC, Rev, TGATCTGAAAAACAGGCA; Pair (3) Fwd, CCGTCCGACATATCATG, Rev, CATTCCCGTCTGAATAATG. The radioactive blot was exposed to a phosphor screen and developed with a Typhoon FLA 9000 phosphor-imager.

## Acknowledgments

We are particularly grateful to Melissa Harrison (University of Wisconsin - Madison) for reading and critiquing early drafts of the manuscript, and to Erika Shor (Center for Discovery and Innovation, Hakensack Meridian Health) and members of the Fox lab for many thoughtful conversations. Support for this work was provided by NIHGM056890 to CAF, a Biotechnology and Biological Sciences Research Council grant BB/N016858 to CAM and CAN, and a Wellcome Trust Investigator Award 110064/Z/15/Z to CAN.

## Supplementary Material

**Figure S1:**
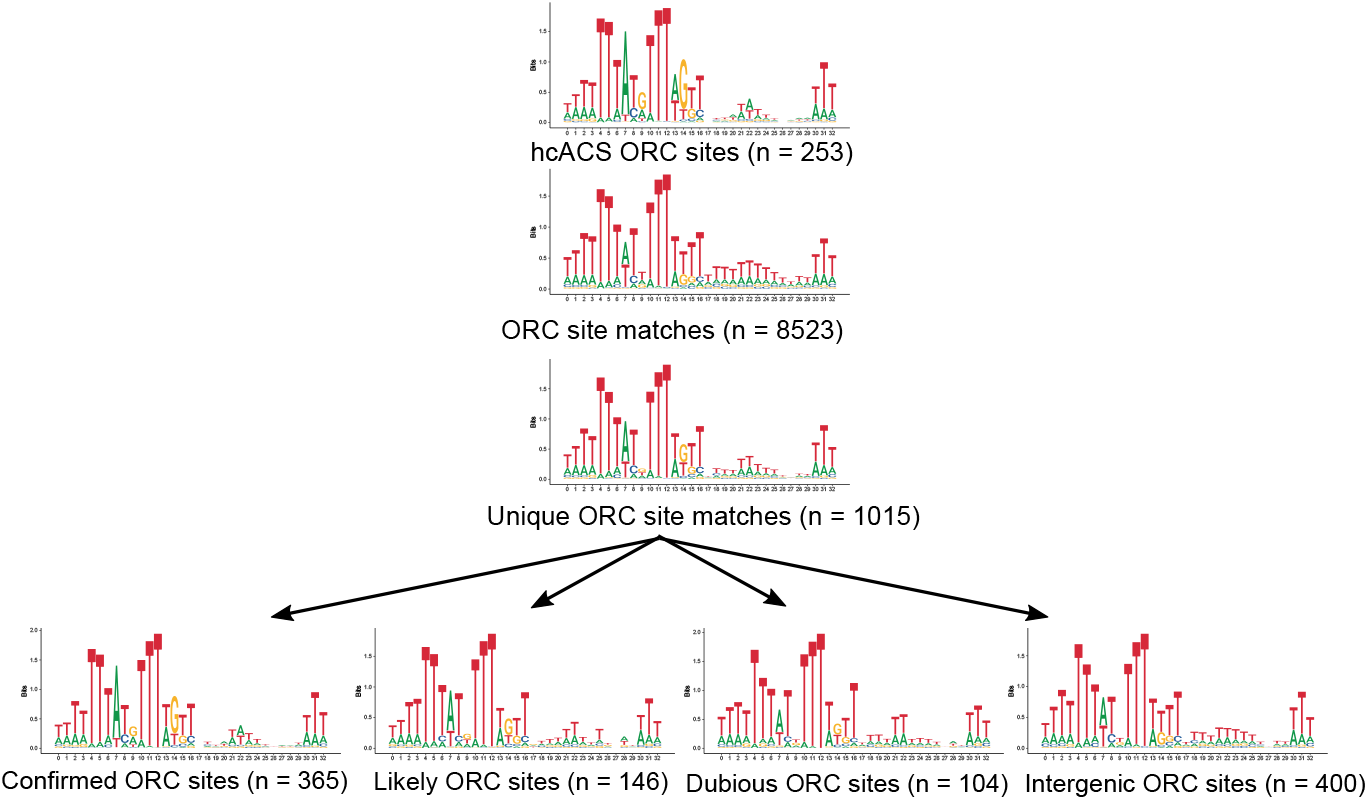
Identification of ORC-binding-site-containing genomic fragments to define the population of ‘origins’ for Z-score analyses. The bioinformatics software Mochiview (57) was used to derive a position specific scoring matrix (PSSM) from all high confidence ORC binding sites (i.e. ORCACS) sequences annotated in (19). The ORC site match collection used here expanded those in the ORCACS to include all likely and dubious origins annotated in OriDB (15). There were 8523 matches meeting −log10 P-values ≥ 4 in the genome (version sacCer3). These matches were mapped to the genome coordinates of all confirmed (n = 410), likely (n = 216) and dubious (n = 203) origins annotated in the OriDB. In addition, matches were mapped to intergenic spaces (n = 6331) annotated in the Saccharomyces Genome Database (SGD). From the intergenic loci that did not overlap OriDB loci (n = 5224 / 6331), we randomly selected 400 non-origin, ORC site containing loci to use in analyses in Figures 2, 3, 5 and 6. When a given locus contained more than one match, the ORC site match with the highest −log10 P value was considered the ORC site match for that locus.

**Figure S2:**
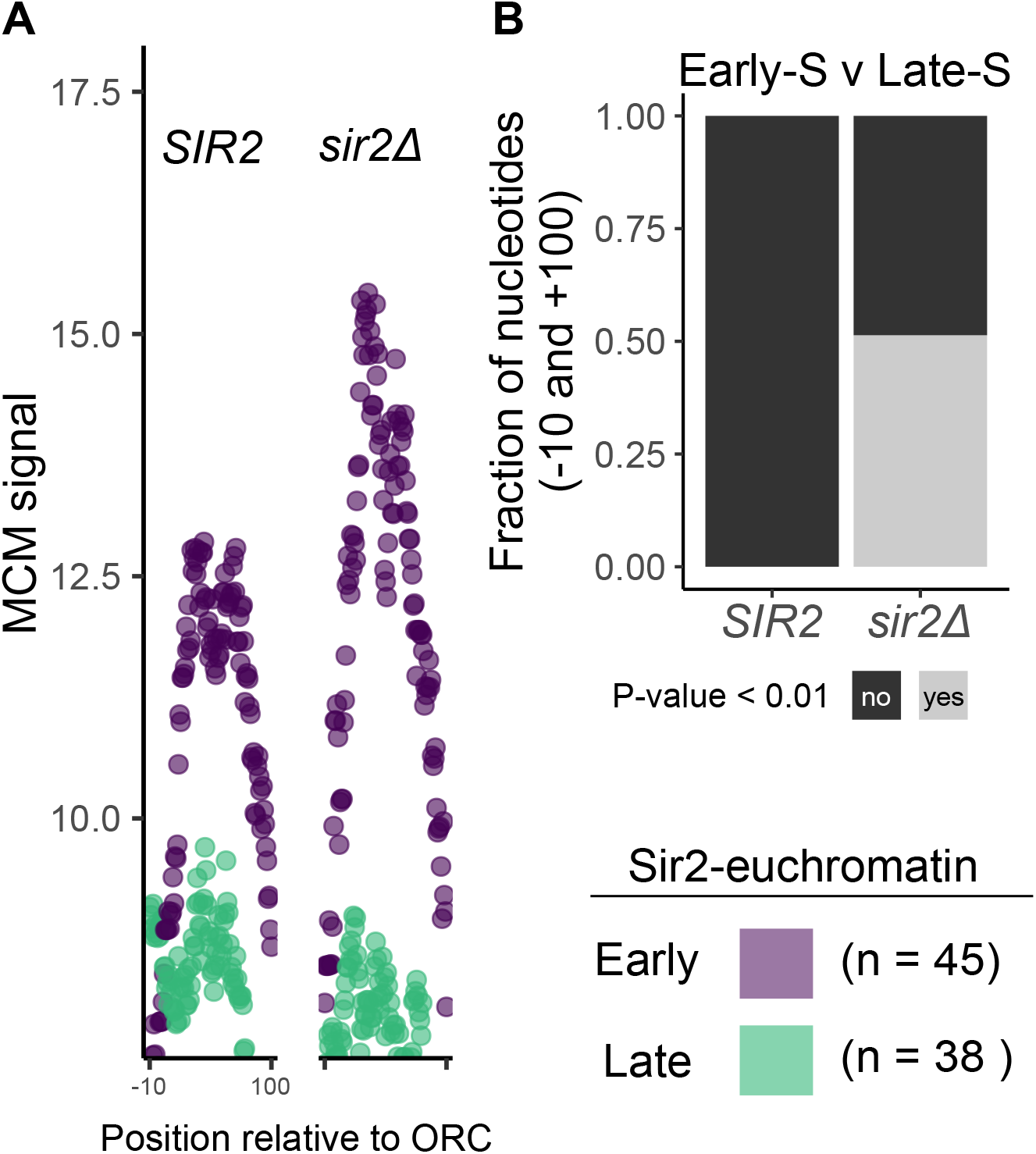
Significant difference in median MCM signal of early and late euchromatic origin cohorts. **A**. Median MCM signals measured in *SIR2* and *sir2*Δ cells within positions −10 through +100 relative to ORC site matches as in **Figure 4C**. **B**. Enrichment of statistically significant differences in MCM medians in *sir2*Δ but not *SIR2* cells. To test whether the differences in the median MCM signals associated with the two cohorts of euchromatic origins reached statistical significance, Wilcoxon rank sum tests were performed on the MCM signal distributions at each position indicated in either genotype. Plotted are the fraction of positions with resulting P-values < 0.01.

**Figure S3:**
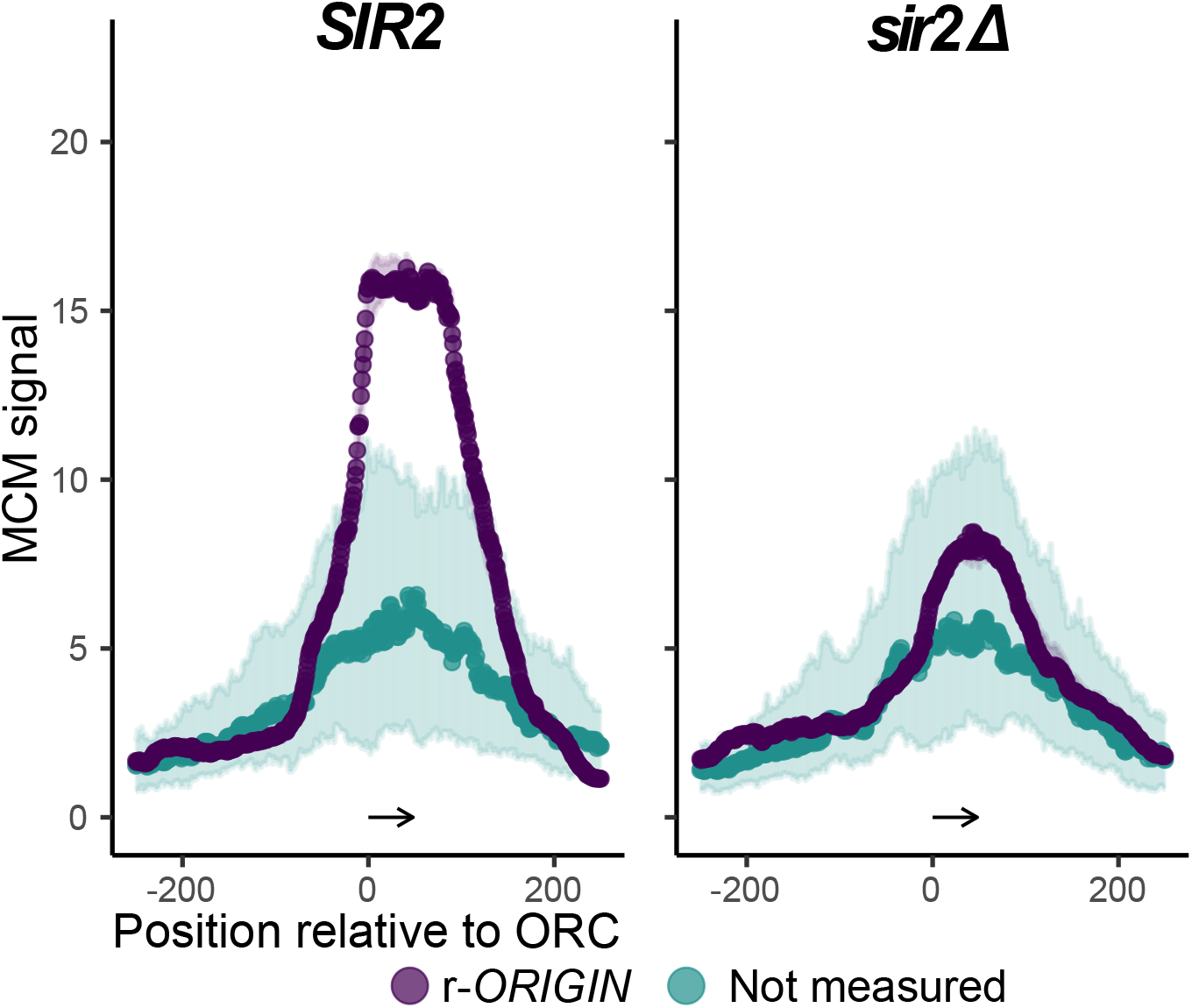
Loss of *SIR2* did not enhance MCM association at the rDNA origin relative to a control origin group. Same approach as in **Figure** 4 only the internally scaled rDNA origin (*r-ORIGIN*) is analyzed and shown in relation to a group of euchromatic origins (Not measured origins, origins lacking Trep values (21)) with little to no change in MCM signal between the two genotypes.

**Figure S4:**
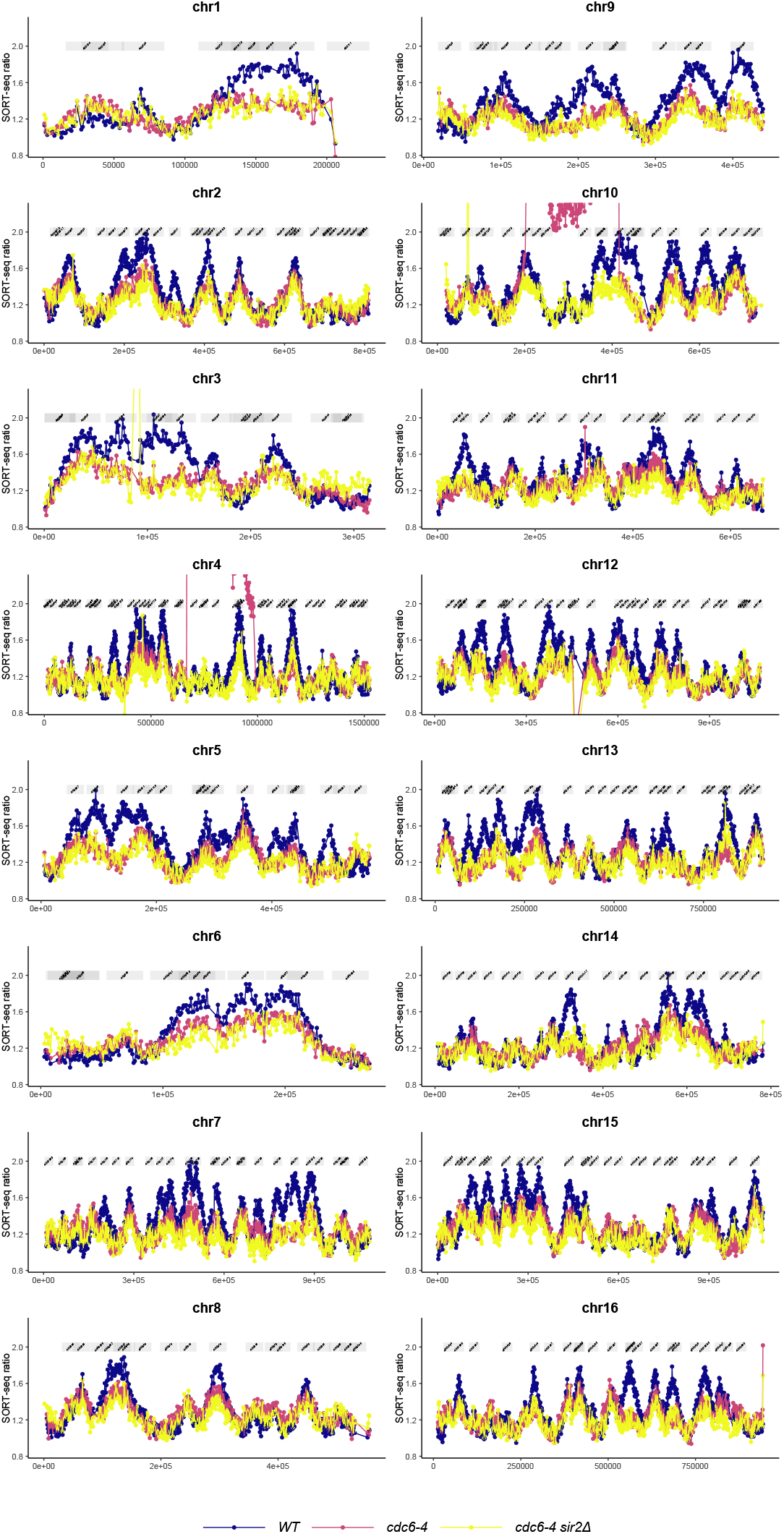
Genome-wide Sort-Seq ratios. The profiles show the relative normalized copy number ratio (y-axis) for each region on all chromosomes (x-axis). The dots represent the binned 1 kb ratios. Profiles are shown for wild-type, *cdc6-4* and *cdc6-4 sir2*Δ cells.

**Figure S5:**
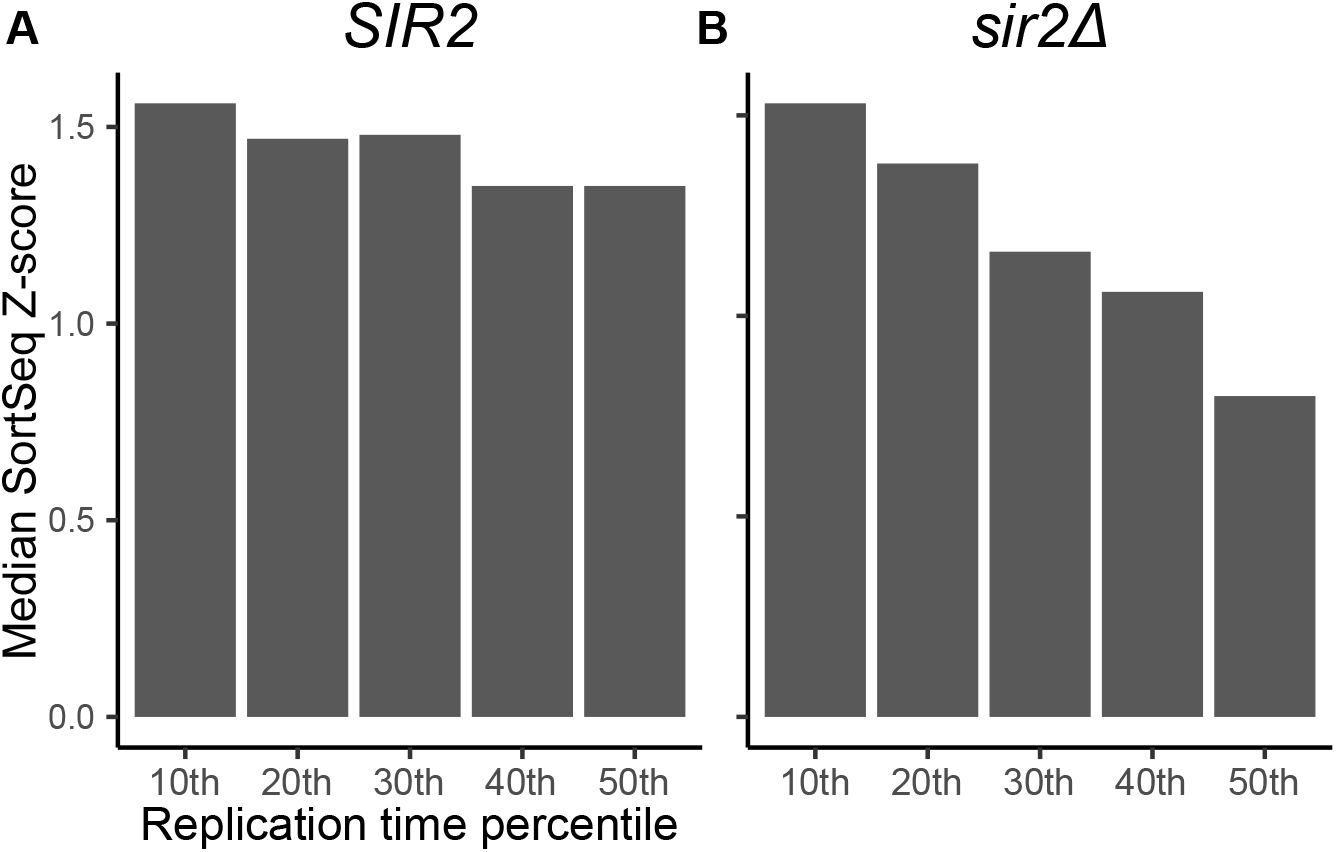
Comparison of SortSeq Z-score medians for euchromatic origins in the top 50th replication time percentiles. **A**. Median Z-scores measured in *SIR2* cells. **B**. Those measured in *sir2*Δ cells. Note the hierarchy of median Z-scores pronounced in *sir2*Δ but not in *SIR2* cells. Consistent with that published previously (30), earlier replicating origins do show a general shift in lower Z-scores in *sir2*Δ cells consistent with delay in replication initiation in S-phase. Specifically, compare the medians measured in *SIR2* and *sir2*Δ for the 30th through the 50th cohorts.

**Figure S6:**
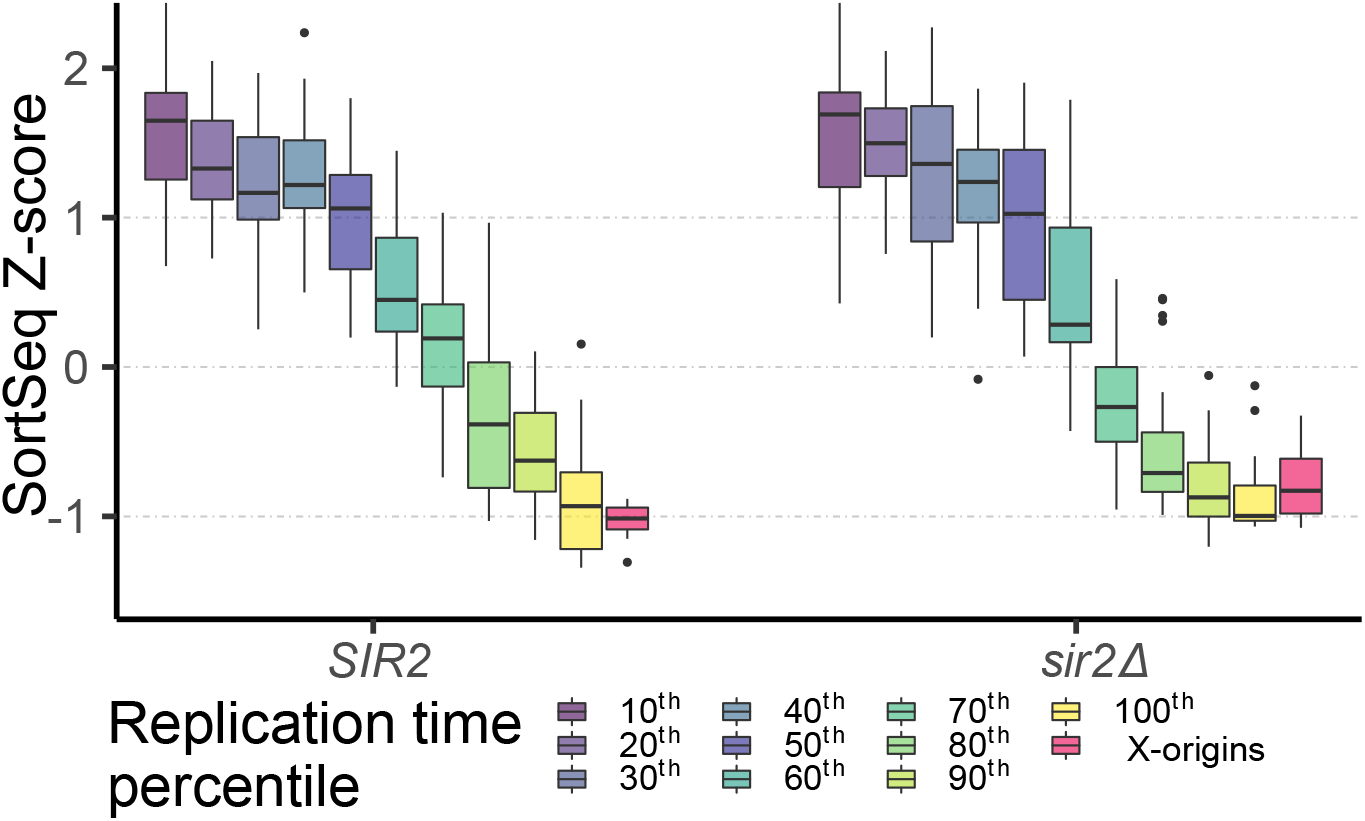
Replication gap observed in an independent data set and in different strains. Analyses of Sort-Seq data from (31) performed as in **Figure** 5**D**.

**Figure S7:**
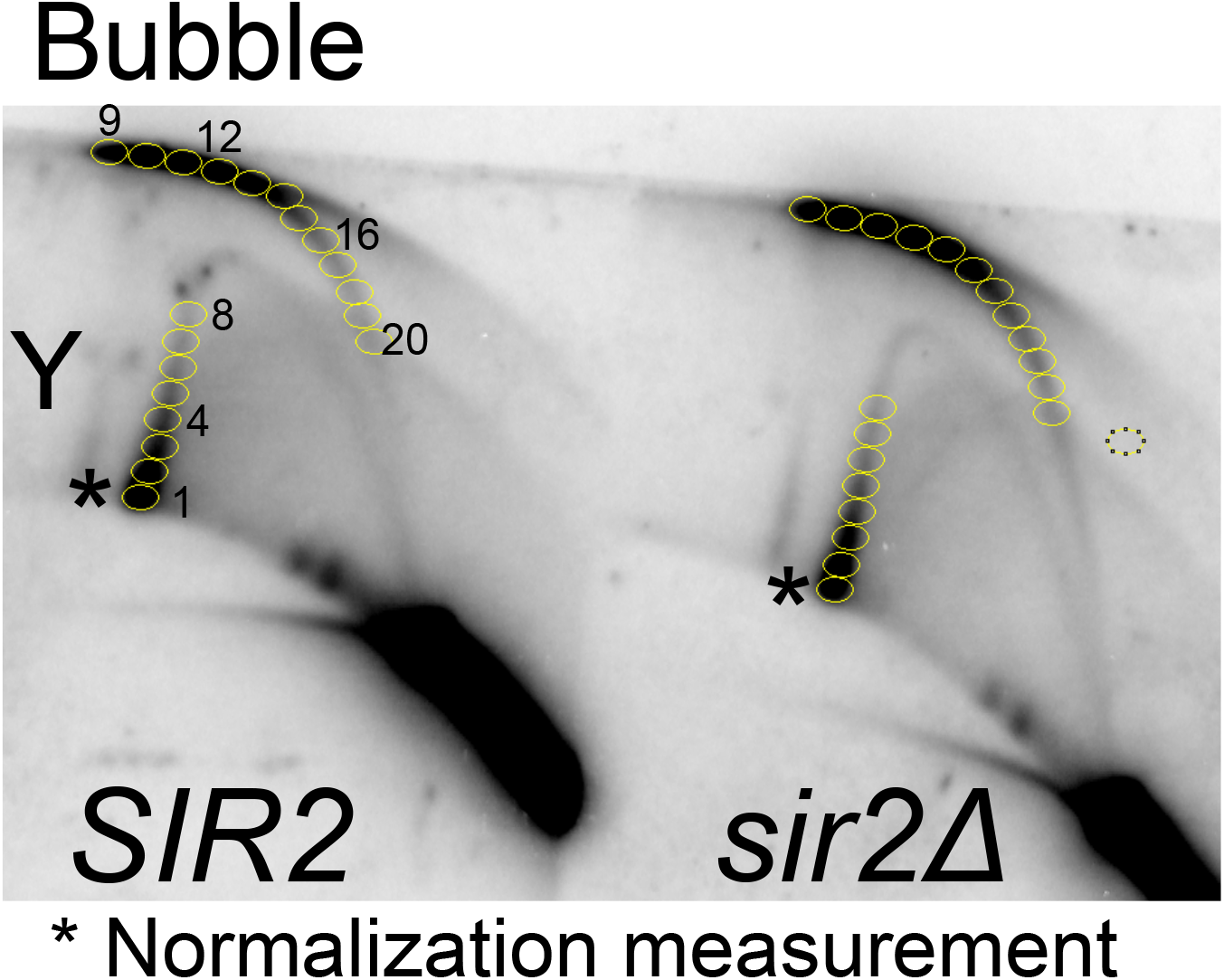
Quantifying signals of replication-intermediates at *ORI922* in *SIR2* and *sir2*Δ? cells. The blot image file generated by the GE Typhoon Phosphorimager was imported into ImageJ 1.51q and processed by measuring the integrated intensities within the indicated circles of identical areas. The * refers to the large fork signal that was used as the denominator for normalizing the signals generated by all the other regions

**Figure S8:**
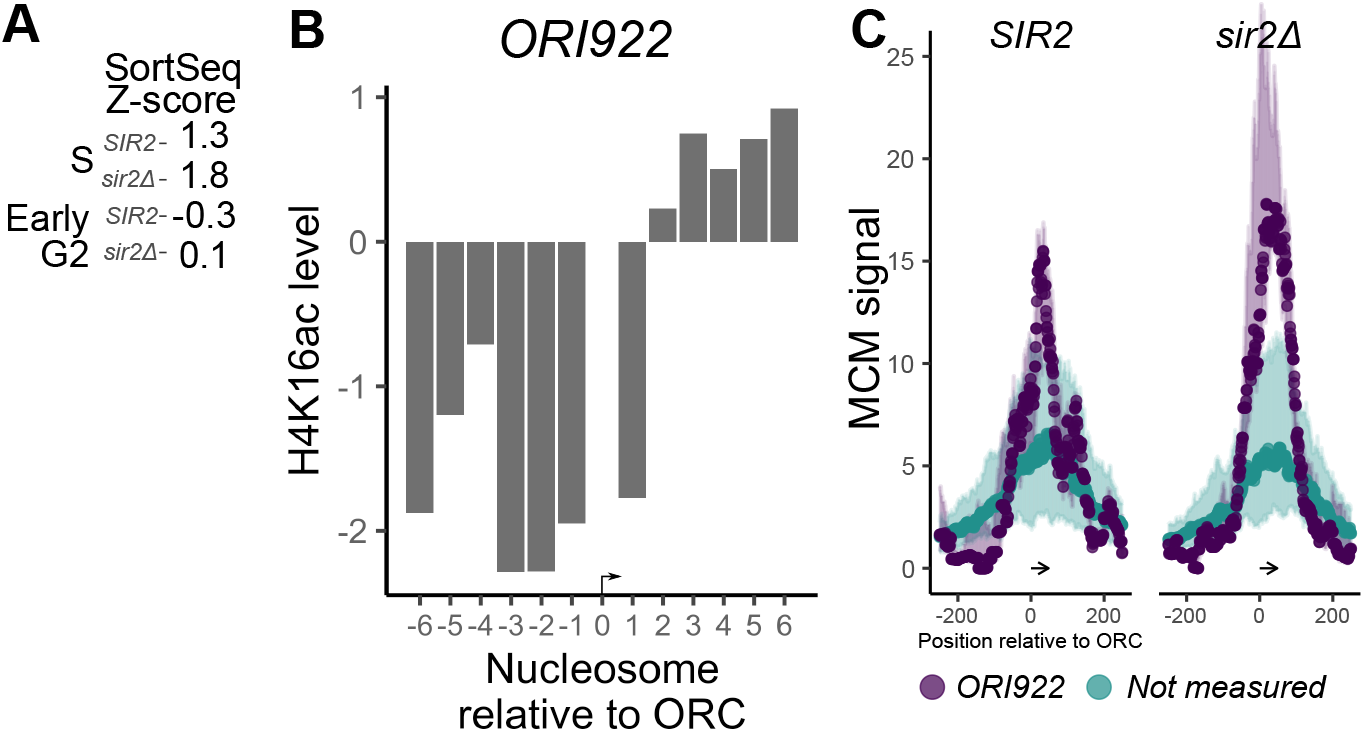
*ORI922* data in this study. **A**. Z-scores assigned to the *ORI922* fragment in indicated cells and cell-cycle phases from data presented in (31). **B**. Normalized H4K16ac levels in *SIR2* cells. **C**. Internally scaled MCM ChIP-Seq data.

